# The Class VIII myosin ATM1 is required for root apical meristem function

**DOI:** 10.1101/2022.11.30.518567

**Authors:** Damilola Olatunji, Natalie M. Clark, Dior R. Kelley

## Abstract

Myosins are evolutionarily conserved motor proteins that interact with actin filaments to regulate organelle transport, cytoplasmic streaming and cell growth. Plant-specific Class XI myosin proteins direct cell division and root organogenesis. However, the roles of plant-specific Class VIII myosin proteins in plant growth and development are less understood. Here, we investigated the function of an auxin-regulated Class VIII myosin, Arabidopsis thaliana Myosin 1 (ATM1), using genetics, transcriptomics, and live cell microscopy. *ATM1* is expressed in the primary root, adventitious roots and throughout lateral root development. ATM1 is a plasma membrane localized protein that is enriched in actively dividing cells in the root apical meristem (RAM). Loss of *ATM1* function results in impaired primary root growth due to decreased RAM size and reduced cell proliferation in a sugar-dependent manner. In *ATM1* loss-of-function roots, columella reporter gene expression is diminished, and fewer columella stem cell divisions occur. In addition, *atm1-1* roots displayed reduced auxin responses and auxin marker gene expression. Complementation of *atm1-1* with a tagged ATM1 driven under the native *ATM1* promoter restored root growth and cell cycle progression in the root meristem. Collectively, these results provide novel evidence that ATM1 functions to influence cell proliferation and columella differentiation in primary roots in response to auxin and sugar cues.

## Introduction

In multicellular organisms, organogenesis relies on the coordination of three interwoven processes: cell proliferation, elongation and differentiation (De Smet and Beeckman, 2011; Harashima and Schnittger, 2010; Stals and Inzé, 2001; Zluhan-Martínez et al., 2021). Cell proliferation events within a developing organ drive growth via an increase in cell number that is triggered and sustained by growth cues, which is a tightly regulated process (Polymenis and Aramayo, 2015). In contrast, cell elongation contributes to cell size regulation, and as a consequence can fine tune organismal body plan (Cnodder et al., 2006; Marshall et al., 2012; Szövényi et al., 2019). During cell differentiation, a variety of cell types constituting an tissue and/or organ become specialized with distinct patterns and forms (Hulskamp et al., 1998). Organogenesis in plants is predominantly post-embryonic and relies on stem cell niches in the shoot and root apical meristems to continuously produce new cells that assume specific differentiation patterns (Burian et al., 2016; Shishkova et al., 2008). Roots are an ideal model to study the coordination of these processes because a growing primary root contains: (i) a meristematic zone, harboring active dividing cells, (ii) a transition zone (TZ), found between the basal meristem and the meristematic zone, containing cells that still have the competence to divide but at relatively slow rate, (iii) an elongation zone (EZ), which has cells with an accelerated expansion rate, mainly in length but not in width and (iv) a differentiation zone (DZ), which has cells that have stopped expansion, but begin to differentiate into specialized tissues. Phytohormones are the major regulator of plant growth responses and a great amount of information has been generated on this in the past decades (Lv et al., 2019; Santner et al., 2009; Shi and Vernoux, 2022), while, the roles of sugars as growth cues are now receiving a great deal of attention. Therefore, understanding the mechanisms of how root organogenesis is regulated in response to both hormonal and metabolic signals is important for furthering our understanding of plant growth and development.

As the by-product of photosynthesis, sugars are translocated to sink organs including roots to orchestrate growth and branching programs, mostly in the form of sucrose, which is subsequently converted to glucose and fructose for the sustenance of energy metabolism (Kaur et al., 2021; Li and Sheen, 2016; Rolland et al., 2006). Genetic and biochemical studies have revealed the link between plant Target of Rapamycin (TOR) and nutrient signaling (Dong et al., 2015; Xiong et al., 2013). Sugar availability is the main activator of TOR kinase in plants resulting in the control of cell proliferation and reprogramming of the transcriptome of the meristems (Xiong et al., 2013). In the presence of glucose, TOR is activated and directly phosphorylates E2F (a/b) transcription factors to maintain the shoot and root meristematic activities (Li et al., 2017). Many of the downstream players in this process are not well understood and it is unknown how stem cell properties are modulated in response to sugars.

Recent studies have revealed the contributions of plant myosins in the regulation of plant growth and developmental programs in response to carbon and hormonal cues (Abu-Abied et al., 2018; Han et al., 2021; Holweg, 2007; Holweg et al., 2003; Ojangu et al., 2018; Olatunji and Kelley, 2020). In plants, myosins are actin-based motor proteins belonging to myosin VIII and XI families. The Arabidopsis genome encodes 13 members of class IX myosins and 4 members of class VIII motor proteins (Haraguchi et al., 2019; Reddy and Day, 2001; Ryan and Nebenführ, 2018). Myosins of the class IX are the most well-studied plant specific myosins to date and their roles have been implicated in rapid cell growth and expansion (Kurth et al., 2017; Peremyslov et al., 2008; Peremyslov et al., 2010). In contrast, information on the roles of class VIII myosins, including ATM1, in plant development still remains limited. Among plant specific myosins, ATM1 was the first member to be identified and sequenced (Knight and Kendrick-Jones, 1993). Immunolocalization studies have shown that ATM1 is localized to the plasmodesmata and new cell plates in Arabidopsis roots (Reichelt et al., 1999). Studies on the full-length or tail domain region of ATM1 fused to GFP expressed under its native promoter indicated that ATM1 is endogenously localized to the plasmodesmata, endoplasmic reticulum, plasma membrane, plastids, and newly formed cell walls (Golomb et al., 2008; Haraguchi et al., 2014). *ATM1* accumulation was also observed in root and shoot apices using a GUS reporter transgene (Haraguchi et al., 2014). ATM1 protein levels are increased in response to exogenous indole-3-acetic acid (IAA) treatment (Kelley et al., 2017). ATM1 plays a key role in sugar-dependent hypocotyl growth, which is driven solely by cell elongation (Olatunji and Kelley, 2020). However, the roles of ATM1 in primary root development are unknown.

Here, we investigated the role of ATM1 in the regulation of root stem cell properties, including proliferation and differentiation. We report that the *atm1-1* mutant exhibits a sugar-dependent short root phenotype due to impaired cell cycle activity. *In situ* DNA labelling and live imaging revealed that ATM1 is required for normal cell proliferation in response to sucrose. Gene expression and auxin response analysis indicate that *atm1-1* root apical meristems exhibit dampened auxin responses in the quiescent center and columella. Transgenic complementation of *atm1-1* mutant with *ATM1pro::GFP-ATM1* or *ATM1pro::GUS-ATM1* cassette can restore root growth and RAM activity. In addition, GFP translational reporters for ATM1 expression *in vivo* indicate that this plasma membrane-localized protein is enriched in root stem cell populations. Altogether these data suggest that ATM1 is required for cell proliferation and differentiation in Arabidopsis roots.

## Results

### *ATM1* promoter activity is high in the root apical meristem

To determine ATM1 expression patterns during root development, stable transgenic lines were generated with *ATM1pro::NLS-GFP-GUS* construct consisting of a 4.5 kb ATM1 promoter sequence. Confocal microcopy of 5-day-old Arabidopsis primary roots (Fig. 1A) and adventitious roots (Fig. 1B) revealed that ATM1 is strongly expressed in apical root cells. Because myosin IX has been implicated in post-embryogenic root formation (Abu-Abied et al., 2018), we then monitored ATM1 expression across the eight stages (Stage I – VIII) of lateral roots (LR) formation (Péret et al., 2009). ATM1 expression was uniformly observed in LR primordia at all stages of LR branching programs (Fig. 1C). Collectively these expression data demonstrate that ATM1 is expressed in developing roots.

**Fig. 1.**
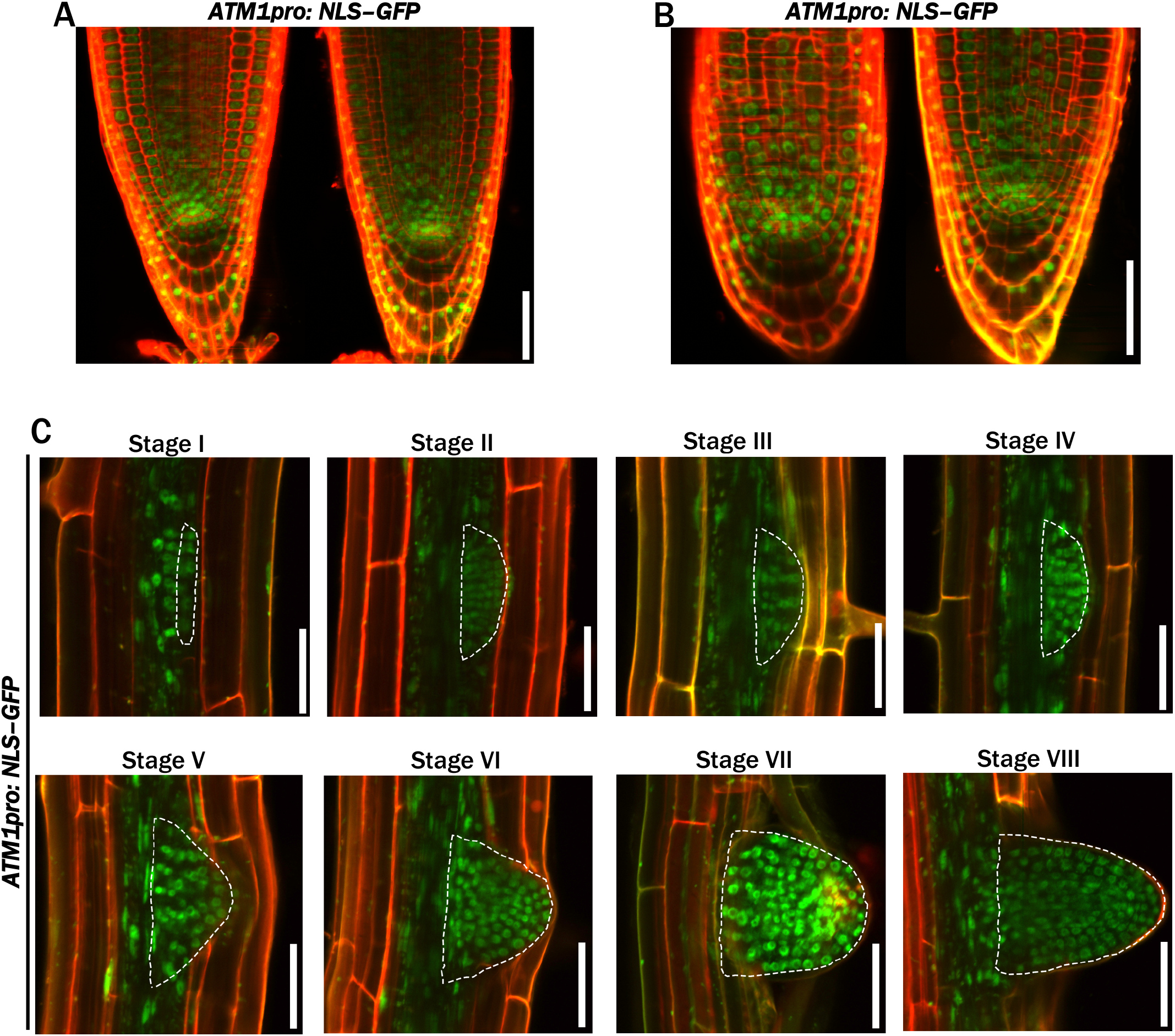
ATM1 promoter is expressed during post-embryogenic root organogenesis in *Arabidopsis thaliana*. (A-B) Transcriptional expression pattern of ATM1 in Arabidopsis roots. *ATM1pro:NLS–GFP* construct expression in the stem cell initials of 5-day-old primary roots (A) and in adventitious roots (B). (C) *ATM1* is expressed in all stages of lateral root development, from initiation (stage 1) to emergence (stage VIII). Dashed white line indicates outline of root development. Scale bars: 50 μm.

### Loss of *ATM1* results in reduced root growth

To examine ATM1 protein expression patterns *in vivo*, stable transgenic lines were generated with *ATM1pro::GFP-ATM1* construct. Confocal microscopy of GFP-ATM1 in Arabidopsis roots revealed protein accumulation at the plasma membrane. In addition, this line showed strong GFP-ATM1 accumulation in the meristematic zone of the RAM and stele initials (Fig. 2A). Collectively these expression data demonstrate that ATM1 is a plasma membrane-localized protein that is present in the primary root.

**Fig. 2.**
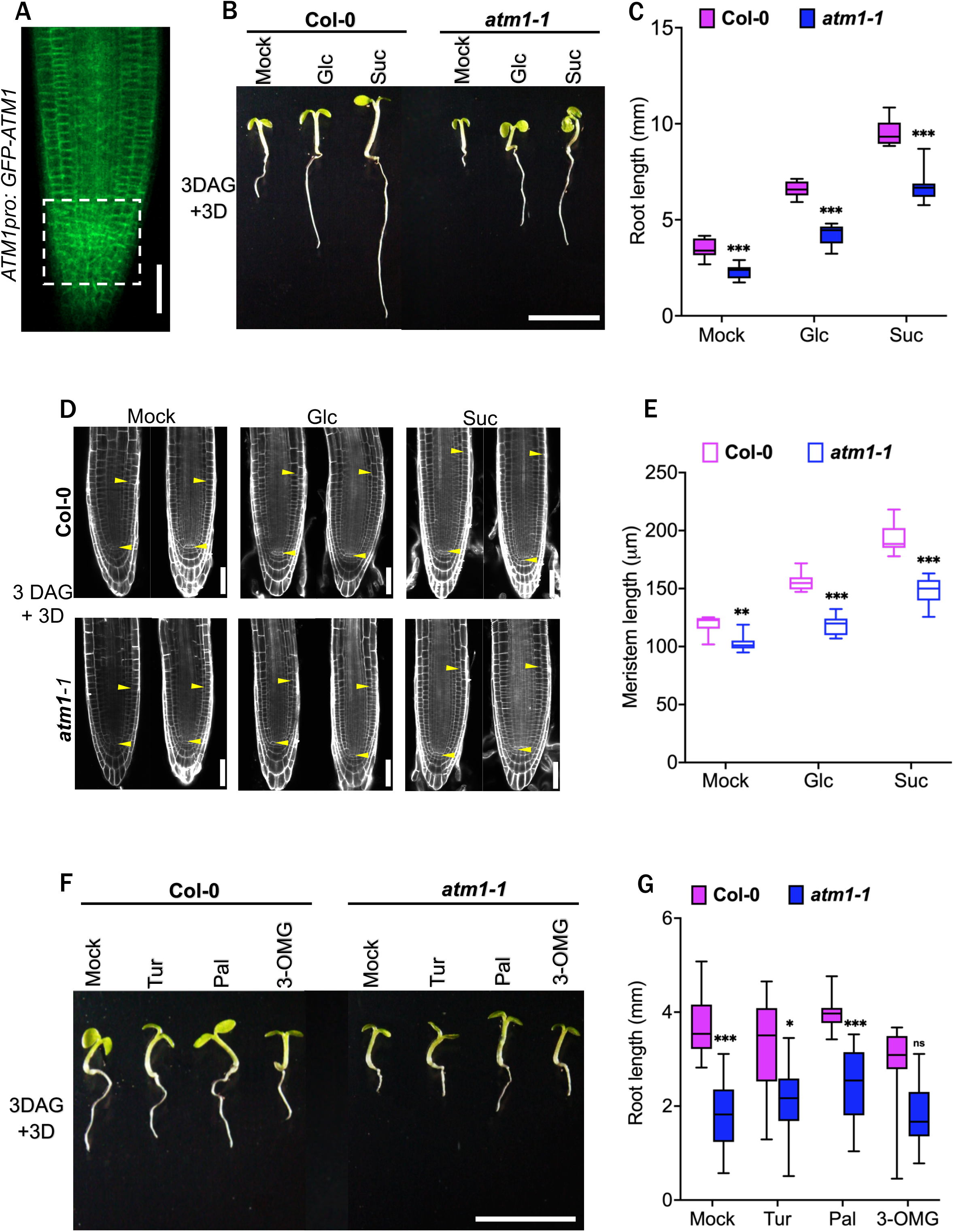
ATM1 a plasma membrane-localised protein required for sugar-activated root growth. (A) GFP-ATM1 protein accumulates at the plasma membrane of the root apical meristem cells in 5-day-old plants. The dashed-square lines represent the stem cell niche with elevated GFP-ATM1 accumulation. (B) Re-activation of root growth in 6-day-old wild-type and *atm1-1* seedlings in response to half strength MS medium (mock) or supplemented with 15 mM Glucose (Glc) or 15 mM Sucrose (Suc) under 15 μmol m-1 s-1, 12-h light/12-h dark conditions. (C) Quantification of root growth across 3 days following Glc or Suc treatment in Col-0 and *atm1-1* seedlings. *n=10*; \*\*\**P*<0.001, two-way ANOVA and Tukey’s multiple comparison test. (D) Confocal images of Col-0 and *atm1-1* roots grown without exogenously applied sugar (mock) or with sugars (Glc or Suc) for 3 days. Roots were stained with propidium iodide. Yellow arrows delineate the root apical meristem (RAM). (E) Root meristem length of 6-day old Col-0 and *atm1-1* plants. *n=10*; \*\**P*<0.01, \*\*\**P*<0.001, two-way ANOVA and Tukey’s multiple comparison test. (F) Phenotypes of 6-day-old Col-0 and *atm1-1* plants grown on 0.5X MS medium with or without 15 mM non metabolizables sugars (Tur: Turanose; Pal: Palatinose and 3-OMG: 3-O-methyl-D-glucose). (G) Root length of Col-0 and *atm1-1* in response to Tur, Pal and 3-OMG. *n=10*; ns, not significant, \**P*<0.05, \*\*\**P*<0.001, two-way ANOVA and Tukey’s multiple comparison test. Box plots extend from 25th to 75th percentile; horizontal lines represent median; whiskers represent minimum to maximum values. Scale bars: 50 μm (A, D), 5 mm (B, F).

To genetically investigate the role of ATM1 in plant root development, we utilized a previously characterized loss-of-function *ATM1* allele (Olatunji and Kelley, 2020), *atm1-1* (SAIL_405_B08). Loss of *ATM1* leads to reduced organ growth in light grown seedlings in the absence of exogenous sugar (Olatunji and Kelley, 2020). Compared to wild-type (WT) Col-0 plants, *atm1-1* root growth was significantly decreased in the absence of exogenous sugar (Fig. 2B,C). Given that ATM1 expression is more pronounced in the region of active cell division in the RAM, we used a previously described root growth re-activation assay (Li et al., 2019; Xiong et al., 2013) to examine how RAM cells respond to sugar signals in *atm1-1*. For this assay, wild-type and *atm1-1* seedlings were grown without sugar under photosynthesis-constrained low light conditions for three days to induce mitotic quiescence. Then, these seedlings were moved to half-strength Murashige & Skoog (MS) medium (mock) or supplemented with glucose or sucrose to re-activate the arrested root meristems. Compared to wild-type Col-0, the *atm1-1* roots have reduced sugar-induced growth (Fig. 2B,C).

The sugar-dependent impairment observed in *atm1-1* roots may be due to altered RAM size. To test this idea, we examined the root meristems of 6-day-old plants grown on control liquid 0.5 x MS medium (mock) or 0.5X MS supplemented with sugars. The *atm1-1* root meristem length was significantly different from that of WT with or without sugar treatment (Fig. 2D,E). Next, we asked whether the reduced-sugar activated root growth in *atm1-1* is attributed to the function of sugar as energy source or signaling molecules. We analyzed WT and *atm1-1* response to several non-metabolizable sugars: a glucose analog (3-O-methylglucose) and two sucrose analogs: palatinose (Pal) and turanose (Tur). Three-day-old quiescent seedlings grown on half-strength Murashige & Skoog (MS) medium were transferred to growth medium supplemented with 15 mM non-metabolizable sugar and scored for activated root growth after 3 days. When compared to WT, none of the glucose or sucrose analog(s) significantly activate the reduced root growth in *atm1-1* (Fig. 2F,G). Taken together, these results suggest ATM1 is essential for sugar-dependent root growth and stem cell activity.

### Complementation of *atm1-1* restored root growth to normal

We then asked if the impaired root growth in the *atm1-1* mutant could be restored. To address this, we cloned the ATM1 full length genomic sequence, tagged with a GFP or GUS reporter protein at the N-terminal driven by ATM1 native promoter (a 4.5 kb fragment upstream of the first annotated ATG in the ATM1 coding sequence), and the generated constructs *ATM1pro::GFP-ATM1* was used for complementation. From the T3 transgenic lines, we performed reverse transcriptase-PCR (RT-PCR) with primers designed to amplify *ATM1* transcripts upstream the site of insertion in SAIL_405_B08 and full-length *ATM1* (Fig. 3A). RT-PCR results showed the expected *ATM1* transcripts in WT, *atm1-1, ATM1pro::GFP-ATM1/atm1-1* plants upstream of the annotated T-DNA insertion site of *atm1-1* (Fig. 3B). Fulllength *ATM1* transcript was not detected in *atm1-1* mutant but was restored to wild-type levels in the transgene containing lines (Fig. 3B). Next, we screened the complemented lines for root growth using five-day old seedlings grown under our growth conditions. On control plates, the root length of the two complemented lines was not significantly different from that of the WT plants (Fig. 3C,D). Notably, sucrose-induced root growth was restored to wild-type levels in *ATM1pro::GFP-ATM1/atm1-1* lines (Fig. 3C,D). Collectively, these results indicate that a functional ATM1 activity is required for root growth.

**Fig. 3.**
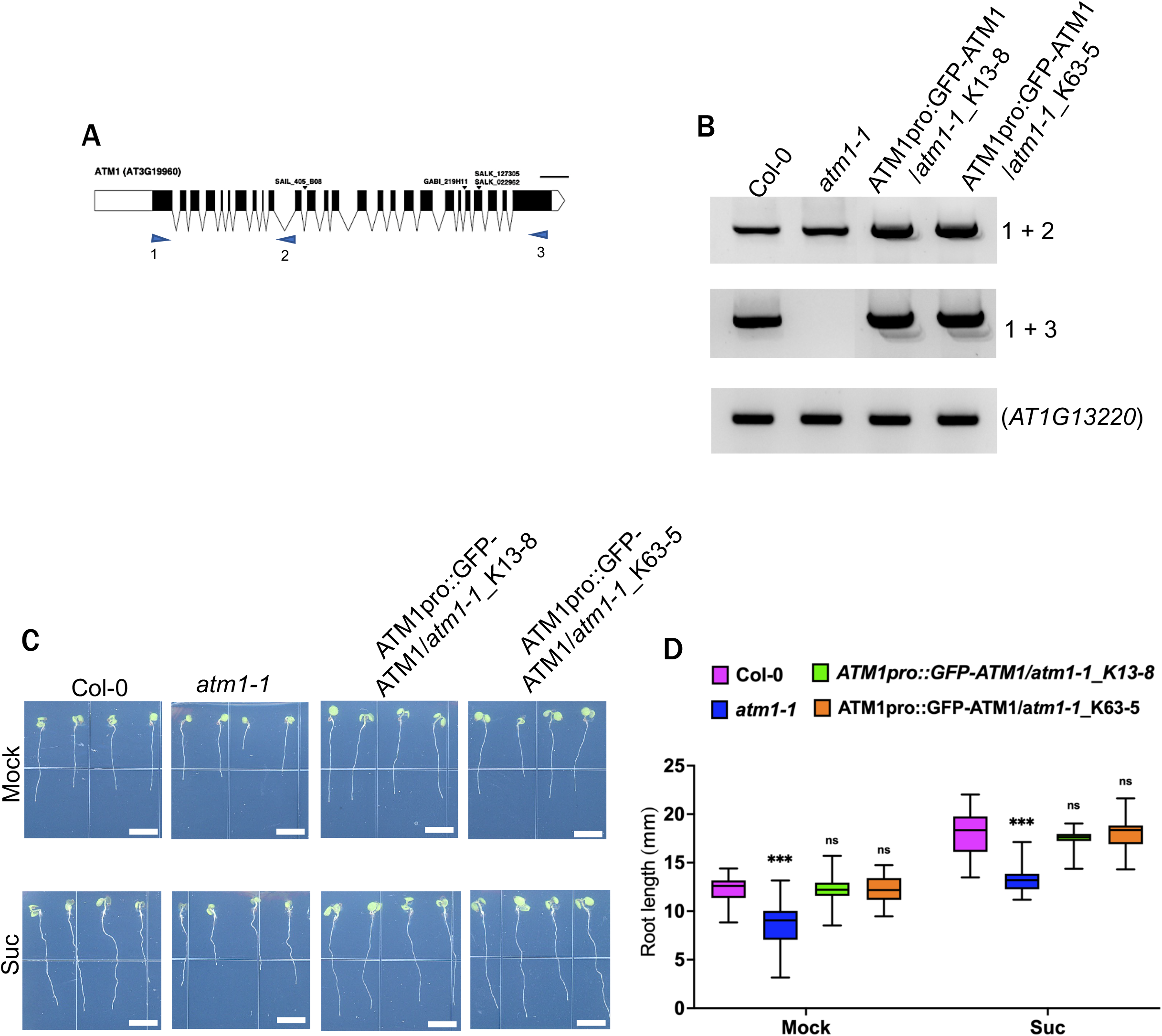
Complementation of *atm1-1* mutant. (A) ATM1 gene model showing the position of SAIL_405_BO8 (*atm1-1*) insertional allele and primers used for PCR amplification. (B) RT-PCR of ATM1 transcripts in CoI-0, *atm1-1* and complemented lines (*ATM1pro: GFP-ATM1/atm1-1_K13-8* or *ATM1pro: GUS-ATM1 /αtm1-1_*K63-5). The top panel shows ATM1 sequences upstream the site of mutation amplified with primers 1 + 2, while the middle panel depicts ATM1 full length produced with primers 1+3. The bottom panel is the transcript level of the endogenous gene (AT1G13220). (C) Root growth in *atm1-1* complemented transgenic lines. Phenotypes of 5-day-old Col-0, *atm1-1and* complemented lines seedlings grown on half strength MS medium (mock) or supplemented with 15 mM Sucrose (Suc) at 45μmol m-1 s-1, 12-h light/12-h dark conditions. (D) Root growth in Col-0, *atm1-1* and the two complemented lines. *n=20*; ns, not significant, ***P<0.001, twoway ANOVA and Tukey’s multiple comparison test. Box plots extend from 25th to 75th percentile; horizontal lines represent median; whiskers represent minimum to maximum values. Scale bars: 5mm.

### Columella cell differentiation is abnormal *atm1-1* roots

To further explore the role of ATM1 in root organogenesis, we examined *atm1-1* roots under normal photosynthetic conditions as previously described (Li et al., 2019). Compared to wild plants, atm*1-1* displayed significantly reduced root growth on sugar-free MS plates (Fig. 4A,B). Moreover, the roots of *atm1-1* seedlings grown on MS medium supplemented with sucrose was significantly decreased compared to wild type plants (Fig. 4A,B). Thus, these data indicate that *ATM1* is required for the regulation of root apical meristem activity. Next, we asked whether the mutation in *ATM1* affects patterning of the root meristem specialized cell types. In Arabidopsis, the root cap comprises two distinct cell types: the lateral root cap (LRC) and columella. Distal to the quiescent center (QC) are the LRC and columella stem cells (CSCs). The CSCs give rise to differentiated columella cells (CCs) containing starch granules required for graviperception (Hong et al., 2015; Su et al., 2017). To investigate whether loss of *ATM1* altered the CSC identity, we crossed the previously characterized *PET111:GFP* enhancer trap line (Clark et al., 2019; Nawy et al., 2005) into *atm1-1*. Typically, *PET111:GFP* marks only the differentiated columella cells (Clark et al., 2019; Nawy et al., 2005). Examination of 5-day-old *atm1-1* roots harboring the *PET111:GFP* transgene revealed significantly diminished expression of this columella marker compared to WT in the presence of sucrose (Fig. 4C,D). Next, we asked whether columella stem cell differentiation in *atm1-1* was impaired by proxy of starch granule presence. To address this question, we stained the roots of 5-day-old WT and *atm1-1* seedlings with Lugol solution. Compared to WT, under sugar-free conditions, the layer of columella stem cell daughter cells (CSCDs) was absent in *atm1-1* (Fig. 4E), suggesting that the competence of these cells to properly differentiate is altered. Under sugar supplementation conditions, the size of the CSCDs and the fully differentiated columella cells (DCCs) were reduced in *atm1-1* compared to WT plants but not the number of cells (Fig. 4E). Despite the mis-expression of the root cap markers in *atm1-1*, surprisingly, GFP expression of the QC marker *WOX5:GFP* is intact the mutant (Fig. S1). These results indicated that ATM1 activity is important for the maintenance of CSC identities.

**Fig. 4.**
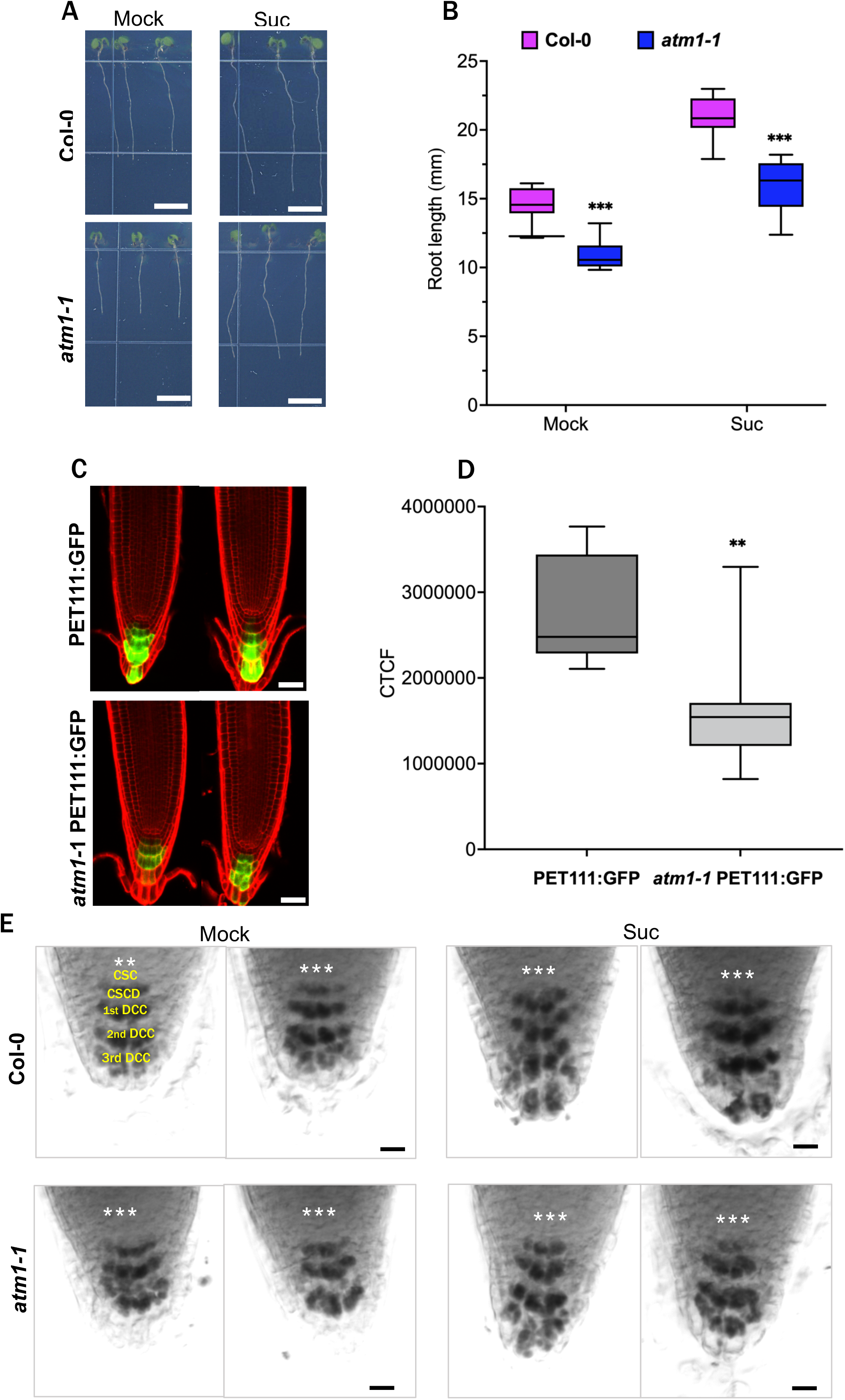
Loss of *ATM1* impacts columella cell differentiation. (A) Phenotypes of 5-day-old *atm1-1* plants with reduced root length compared to wild type on 0.5 X Murashige and Skoog (MS) medium without or with 15 mM Sucrose (Suc) at 45 μmol m-1 s-1 LD conditions. (B) Quantitative analysis of root growth in Col-0 and *atm1-1* seedlings. *n=10*; \*\*\**P*<0.001, two-way ANOVA and Tukey’s multiple comparison test. (C) Examination of columella cells marker (PET111:GFP) accumulation in wild type and *atm1* roots germinated on 0.5 X Murashige and Skoog (MS) medium with 15 mM Sucrose (Suc). (D) Computed average florescence intensity in PET111:GFP and *atm1-1* PET111:GFP. *n=7-8*; \*\**P*<0.01 (unpaired two-tailed Student’s t-test). (E) Starch granule accumulation in Col-0 and *atm1-1* columella cells. Roots of 5-day-old wild type and *atm1-1* plants were stained with Lugol solution prior to imaging. Abbreviations used: Quiescent center: ***; CSC: columella stem cells; CSCD: columella stem cell daughters; 1st DCC:1st layer of differentiated columella root cap cells; 2nd DCC: 2nd layer of differentiated columella root cap cells; 3rd DCC: 3rd layer of fully differentiated columella root cap cells. Box plots extend from 25th to 75th percentile; horizontal lines represent median; whiskers represent minimum to maximum values. Scale bars; 5 mm (A), 40 μm (C), 50 μm (E).

### Transcriptomic analysis of *atm1-1* seedlings in response to sugars

To discover the molecular mechanism controlled by *ATM1* during plant development in response to sugar signaling, we performed bulk RNA-seq analysis using whole seedlings grown with and without exogenously applied sugars (glucose and sucrose) (Table S1). From the transcriptome data, we identified differentially expressed genes (DEGs) that are specifically modulated by sugar molecules in both Col-0 and *atm1-1* mutant compared to mock treatment with significance of *q*-value ≤ 0.1. Previous studies have examined the effects of sucrose and glucose on transcription in Arabidopsis (Mishra et al., 2009; Shulse et al., 2019). In order to uncover sugar-dependent gene regulation that was specifically altered in *atm1-1*, we examined these DEGs in more detail. In total, 32 up-regulated DEGs were observed in *atm1-1* compared to WT under mock conditions, while *atm1-1* seedlings treated with glucose and sucrose relative to mock samples had 406 and 1488 DEGs respectively (Fig. 5A-C (right), Table S1). Among these samples, 1154 down-regulated DEGs were obtained in *atm1-1* sucrose vs mock samples, followed by glucose-treated *atm1-1* seedlings compared to mock samples (134), and control medium-treated *atm1-1/WT* plants had the lowest number (87) of repressed DEGs (Fig. 5A-C (left), Table S1).

**Fig. 5.**
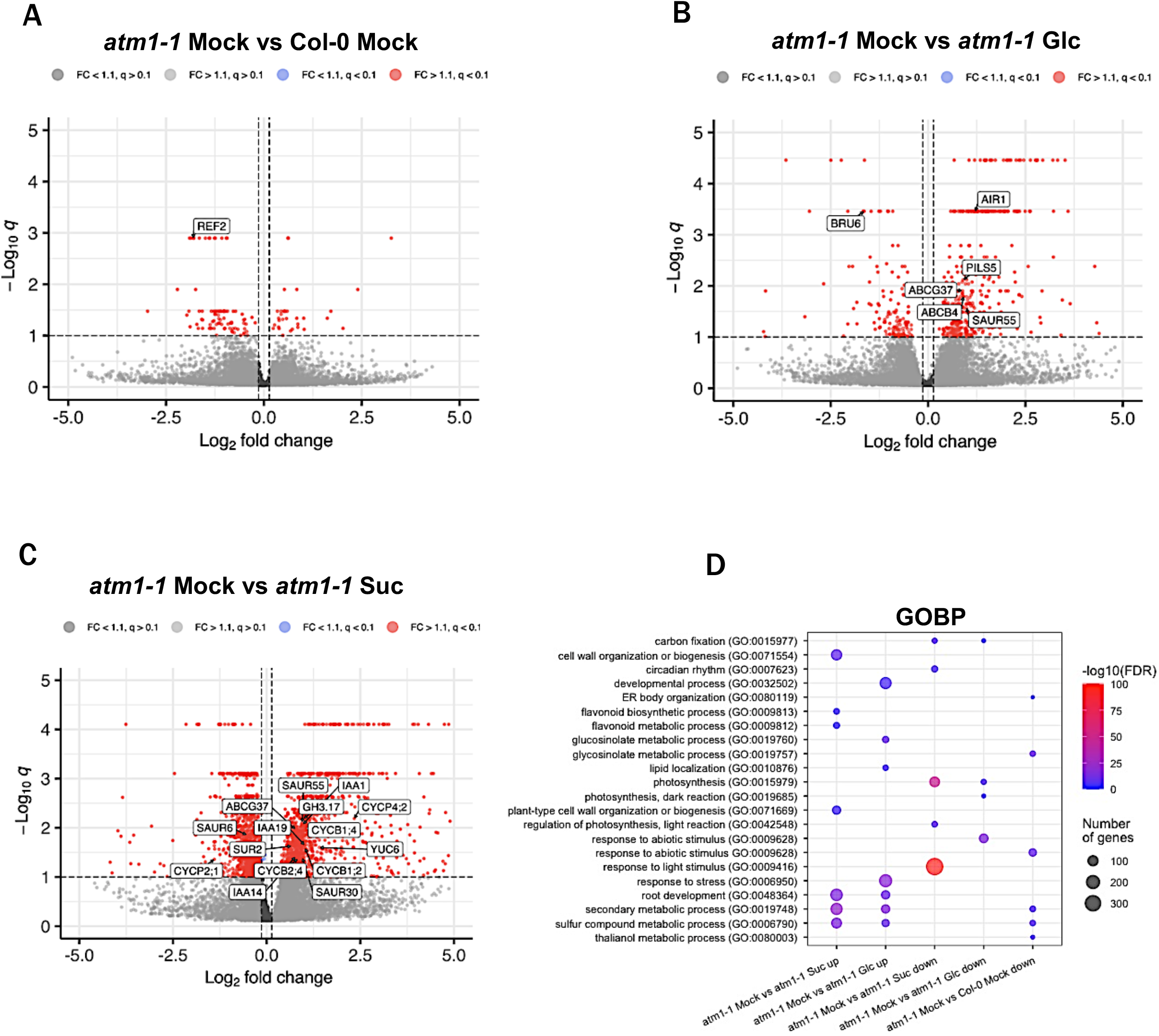
Transcriptomic analysis of *atm1-1* revealed key biological processes. (A-C) Volcano plots showing candidate DEGs up-and-down regulated in *atm1-1* vs Col-0 under Mock treatment (A), in glucose treated *atm1-1* plants (B), and under sucrose treatment compared to mock conditions (C). The repressed and up-activated DEGs are on the left and right sides respectively of the Volcano plots. (D) Enriched gene ontology of biological processes (GOBP) among up-and-down-regulated genes in *atm1-1* under different treatment conditions. The colour depicts significance of the enrichment (-log10(FDR)). The circles represent the number of genes linked to a specific GO term.

Sucrose-regulated auxin pathway genes in both wild-type Col-0 and *atm1-1* include numerous *SMALL AUXIN UPREGULATED* (*SAUR*) genes, *AUXIN-INDUCED IN ROOT CULTURES 1* (*AIR1*), *AIR3, PLEIOTROPIC DRG RESISTANCE 9/PIS1/ABCG37*, and *YADOKARL* (*YDK1*) (Table S1). Notably, in *atm1-1* several classical auxin marker genes are upregulated including *YUCCA3, YUC6, YUC8, GRETCHEN HAGEN 3.17* (*GH3.17*), *IAA19* and *IAA29* (Fig. 5C, Table S1). These genes are not auxin-upregulated in Col-0, suggesting that these gene expression events are due to specific sucrose dependent cues in the absence of *ATM1*. In addition, these data also suggest that sucrose can both induce and repress many key auxin pathway genes, which is consistent with a previous study (Mishra et al., 2009).

Next, to understand the major molecular processes that are significantly enriched in *atm1-1* upon sugars treatment, we performed gene ontology analysis. No significantly enriched GO terms were identified for control medium-treated *atm1-1*/WT plants DEGs, but GO terms such as root development, secondary metabolic process and sulfur compound metabolic process were enriched in sucrose and glucose-treated *atm1-1* samples compared to mock-treated *atm1-1* seedlings, suggesting the essential role of sugar signaling in ATM1-mediated developmental processes (Fig. 5D). Among the enriched GO terms in the down-regulated DEGs for *atm1-1* plants fed with sugar molecules are carbon fixation, photosynthesis, and response to light stimulus. These results suggest that besides exogenously applied sugars, light-mediated photosynthetic processes are essential for hormones biosynthesis and as carbon sources during ATM1-controlled.

### ATM1 activity is dependent on auxin-sugar signaling

Auxin is a central regulator of root growth, and the roles of multiple auxin biosynthesis, transport and signaling pathways have been implicated in *de novo* root organogenesis in plants (Casimiro et al., 2001; Lavenus et al., 2013; Mishra et al., 2022; Olatunji et al., 2017; Roychoudhry and Kepinski, 2022; Singh et al., 2020). Within the primary root, auxin levels in the QC and columella are associated with columella differentiation (Brumos et al., 2018; Ding and Friml, 2010). Because the transcriptomic analysis of *atm1-1* indicated sucrose-activated gene expression of auxin biosynthesis, metabolism, and signaling genes (Fig. 5C, Table S1), and columella marker expression and divisions are altered in *atm1-1* (Fig. 4C,D), we asked if loss of *ATM1* can impact auxin signaling in the RAM. To examine this *in vivo*, we crossed the previously described auxin response marker *DR5:GFP* (Friml et al., 2003) into *atm1-1*. Confocal imaging of *DR5:GFP* in 5-day-old *atm1-1* roots revealed a significant reduction in auxin response with and without exogenous sugar application compared to WT (Fig. 6A,B), suggesting a down regulation of auxin signaling in the mutant under the experimental conditions tested. Collectively, these data indicate that the spatiotemporal regulation of auxin pathways is impaired in the absence of *ATM1* activity and this response is linked to sugar during root development.

**Fig. 6.**
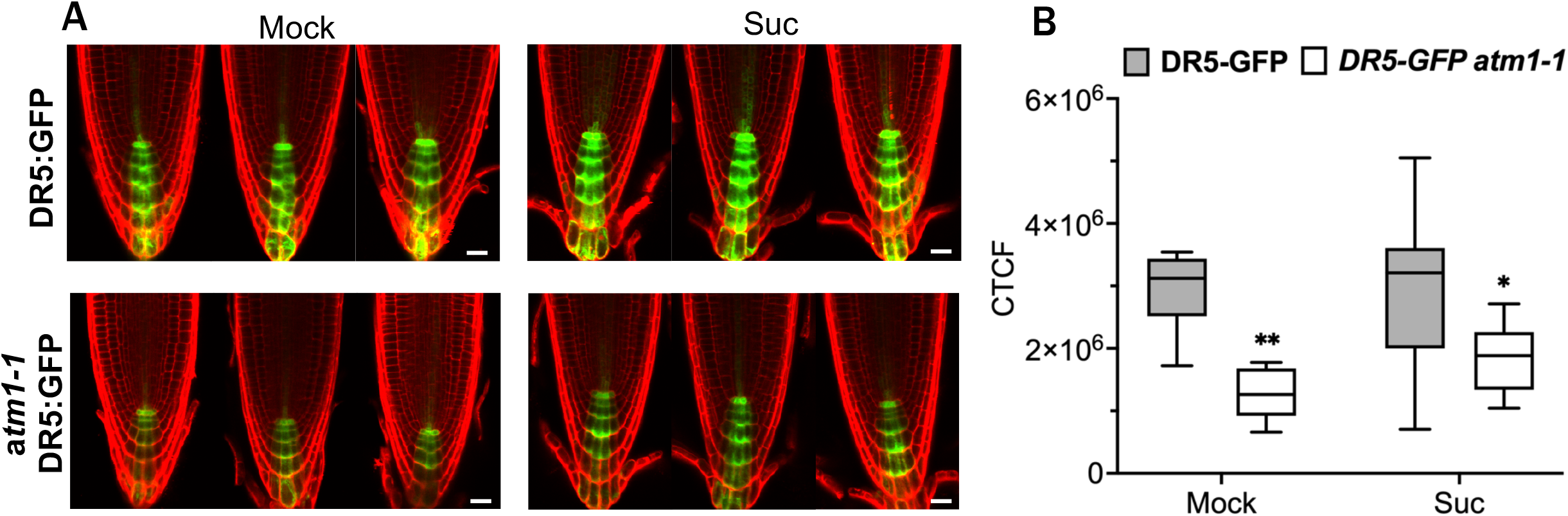
*atm1-1* mutation alters auxin responses *in planta*. (A) Expression of auxin response marker DR5-GFP in *atm1-1* mutant. Roots of 5-day-old seedlings grown on 0.5 X Murashige and Skoog (MS) medium without or with 15 mM Sucrose (Suc) were counterstained with propidium Iodide (PI). (B) Calculated corrected total cell fluorescence in DR5:GFP and *atm1-1* DR5:GFP plants. *n=8-10*; \**P*<0.05, \*\**P*<0.01, two-way ANOVA and Tukey’s multiple comparison test. Box plots extend from 25th to 75th percentile; horizontal lines represent median; whiskers represent minimum to maximum values. Scale bars: 50 μm.

### Cell cycle activation is impaired in *atm1* roots

In addition to auxin marker genes being differentially expressed in *atm1-1*, many cell cycle genes were also induced in *atm1-1* in response to sucrose treatment (Fig. 5C), including B-type *CYCLIN* genes. To verify if cell cycle regulation is influenced by ATM1, we deployed nucleosides analogue 5-ethynyl-2′-deoxy uridine (EdU) to label newly replicating DNA in the root meristems. Nuclear EdU staining is an ideal protocol to obtain information on S-phase cells within developing tissues (Echevarría et al., 2021). For effective investigation of cell cycle progression in *atm1-1* roots, we used three-day-old mitotic quiescent seedlings grown under photosynthesis-constrained light conditions. These quiescent seedlings were treated with MS medium or with either glucose or sucrose for 24 hours prior to *in situ* labelling of the roots with EdU. Under mock treatment, the EdU-labeled DNA in *atm1-1* was not significantly different from WT (Fig. 7A,B). Upon sugar exposure, the quantity of stained cells entry the S phase of cell cycle was significantly reduced in *atm1-1* compared to WT (Fig. 7A,B).

**Fig. 7.**
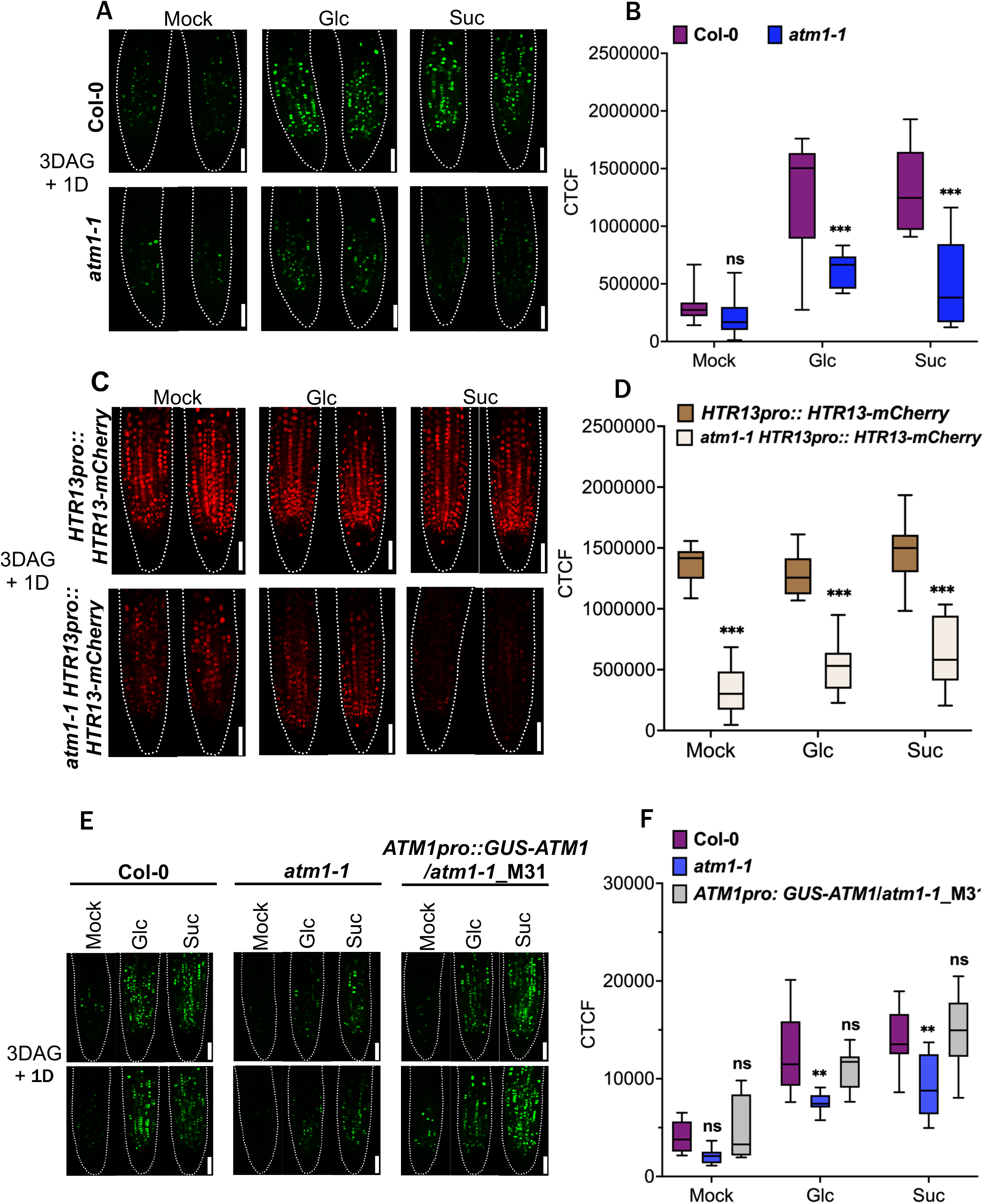
*atm1-1* is defective in S-phase entry of cell cycle. (A) EdU staining reveals reduced number of proliferation cells in *atm1-1* mutant. 3-day-old quiescent Col-0 and *atm1-1* seedlings were treated with half strength MS medium (mock) or supplemented with either 15 mM Glucose (Glc) or 15 mM Sucrose (Suc) and grown for 1 additional day at 15μmol m-1 s-1, 12-h light/12-h dark conditions prior to EdU staining *in situ*. Dashed line represents outline of the root. (B) Quantified average corrected total cell fluorescence (CTCF) in Col-0 and *atm1-1* under different treatment conditions. *n=10*; ns: not significant, \*\*\**P*<0.001, two-way ANOVA and Tukey’s multiple comparison test. (C) A component of Plant Cell Cycle indicator (PlaCCI), *HTR13pro:: HTR13-mCherry*, shows diminished expression in *atm1-1* mutant. The level of the cell proliferation marker H3.1-mCherry expression was reduced in *atm1*. Dashed line depicts outline of the root. (D) Mean total mCherry fluorescence in plants expressing *HTR13pro:: HTR13-mCherry. n=9-10*; \*\*\**P*<0.001, two-way ANOVA and Tukey’s multiple comparison test. (E) S-phase entry of cell cycle is restored in *ATM1pro:GUS-ATM1/atm1-1* as shown by EdU staining signals *in situ*. (F) Computed average corrected total cell fluorescence (CTCF) in Col-0, *atm1-1* and *ATM1pro:GUS-ATM1/atm1-1* under different treatment conditions. Data are presented as means with box and whisker plot. *n=10*; ns, not significant, \*\**P*<0.01, two-way ANOVA and Tukey’s multiple comparison test. Box plots extend from 25th to 75th percentile; horizontal lines represent median; whiskers represent minimum to maximum values. Scale bars: 50 μm (A, C, E).

To further substantiate the role of ATM1 in control of cell cycle events, we crossed the previously described Plant Cell Cycle Indicator (PlaCCI) marker (Desvoyes et al., 2020) into *atm1-1*. The PlaCCI transgene contains three fluorescent markers to monitor cell cycle status, including *HTR13pro::HTR13-mCherry* that reveals cells in S- or G2-phase of the cell cycle. Compared to WT plants carrying *HTR13pro::HTR13-mCherry*, the expression of the HTR13-mCherry signal was significantly reduced in *atm1-1* RAM cells (Fig. 7C,D). We further examined the root meristem of the complemented lines to determine if the competence of the *atm1-1* roots to undergo DNA replication is restored. Our results showed that the root meristem of the quiescent *ATM1pro: GUS-ATM1/atm1-1* seedlings labeled with EdU stain has acquired the capacity to activate S phase dynamics of the cycle when compared to WT plants root meristem (Fig. 7E-F). Taken together, these results suggest ATM1 plays a role in cell proliferation in the root meristem.

## Discussion

In the past years, tremendous progress has been made in plant specific myosin research, particularly in the area of auxin and sugar-regulated developmental processes (Abu-Abied et al., 2018; Han et al., 2021; Holweg, 2007; Holweg et al., 2003; Ojangu et al., 2018; Olatunji and Kelley, 2020). Spatiotemporal studies on myosin expression patterns have revealed the accumulation of myosin XI and VIII in numerous tissues such as flowers, inflorescences, stems, siliques, and roots, suggesting that myosins are required for development at the whole plant level (Haraguchi et al., 2014; Park and Nebenführ, 2013). The use of Arabidopsis root system has extensively aided our knowledge of the molecular and cellular mechanism controlling root development from initiation, patterning to emergence, wherein many key and novel genes functions have been identified (Casimiro et al., 2003; Celenza et al., 1995; Péret et al., 2009; Torres-Martínez et al., 2022). In this study, we investigated how an auxin-regulated Class VIII myosin, ATM1, controls plant root development. Here, a GFP transcriptional reporter indicates that *ATM1* is expressed in primary roots (Fig. 1A) as previously reported (Clark et al., 2019; Haraguchi et al., 2014) and in secondary roots (Fig. 1B,C), indicating that *ATM1* may play a role in root development. Furthermore, *ATM1* expression occurs in a gradient along the RAM that is reminiscent of the *PLETHORA* genes, which are known to positively regulate RAM renewal in response to auxin cues (Mähönen et al., 2014; Santuari et al., 2016). However, ATM1 is not known to be a target of PLT transcription factors (Santuari et al., 2016). Other transcription factor(s) may be important regulators of *ATM1* expression in the RAM, which may be determined in future studies.

Myosin proteins are known to have complex subcellular localization patterns that can include cytoskeletal associations with actin and/or microtubules, plasmodesmal and/or plasma membrane localizations, or at newly formed cell plate (Golomb et al., 2008; Kurth et al., 2017; Nebenführ and Dixit, 2018; Peremyslov et al., 2010; Ryan and Nebenführ, 2018). Myosins can also be associated with vesicular trafficking and move cargo within the cell (Perico and Sparkes, 2018; Ryan and Nebenführ, 2018; Titus, 2018). Here, we confirm that ATM1 is a plasma-localized protein *in vivo*, with strong accumulation in the stem cell niche of the RAM (Fig. 2A). This data suggests that ATM1 may function at the membrane to influence cellular properties within developing roots.

Root morphogenesis is generally activated by sugar molecules (Li et al., 2017; Xiong et al., 2013) generated in the shoot via photosynthesis. To investigate links between photosynthetically derived sugars and the *atm1-1* short root phenotype we used previously described low light conditions (Li et al., 2019). Under depleting endogenous sugar conditions, the short RAM phenotype in *atm1-1* could be attributed to reduced uptake of exogenously applied sugar by the roots (Fig. 2B-E). The use of non-metabolizable analogues of glucose and sucrose have been widely used to understand the role of sugar-mediated signaling pathway in plants (Cortes et al., 2003; Fernie et al., 2001; Gonzali et al., 2005). In addition, turanose insensitivity is associated with altered auxin homeostasis and altered expression of *WOX5*, a central organizer of the quiescent center (Gonzali et al., 2005). To investigate the response of *atm1-1* roots to non-metabolizable sugars, the mutant and WT Col-0 were exposed to turanose, palatinose and 3-OMG. None of these non-metabolizable sugars significantly reactivated *atm1-1* root growth as compared to WT. However, we cannot rule out the fact that the non-metabolizable sucrose isomer palatinose may act as a signaling molecule(Ramon et al., 2008), since they slightly stimulate shoot and root growth in *atm1-1* (Fig. 2F).

Moreover, the fact that *atm1-1* showed impaired root growth under normal photosynthetic conditions (Fig. 3C,D) corroborates the idea that the short root phenotype is due to defects in shoot-to-root sugar transport. In addition, the *atm1-1* short root phenotype can be complemented by a *ATM1pro::GFP-ATM1* transgene (Fig. 3B-D), which opens the door for future biochemical and imaging studies with this fluorescently tagged version of ATM1 expressed under a native promoter.

Studies have shown that altered sugar metabolism can cause a delay in distal stem cell differentiation (Pignocchi et al., 2021; Wang et al., 2018). Furthermore, exogenous application of sugars has been reported to restore delayed distal stem cell in several mutants suggesting that there is a common regulatory pathway underpinning this process (Pignocchi et al., 2021; Racolta et al., 2014; Wang et al., 2018). Therefore, the diminished expression of the columella cell marker *PET111:GFP* in *atm1-1* grown in the presence of sucrose indicates that ATM1 modulates columella stem cell initial formation via a sugar-dependent pathway (Fig. 4C-D). In addition, we showed that sugar treatment (glucose or sucrose) completely reinstates the number of differentiated columella stem cells in *atm1-1* mutant, but not their cell size (Fig. 4E). Based on these findings, we propose that ATM1 plays an essential role in the proper division and maintenance of columella stem cell initials via a sugar-mediated pathway.

The interplay between sugar and auxin has been shown to influence diverse aspects of plant development including root architecture and growth (Mishra et al., 2009; Mishra et al., 2022). Many Previous work has demonstrated that exogenous application of sugars promotes auxin biosynthesis and auxin-regulated gene expression (Mishra et al., 2009; Mishra et al., 2022; Sairanen et al., 2012) Here, the examination of glucose and sucrose regulated gene expression in wild-type and *atm1-1* seedlings revealed the downregulation of genes involved in auxin signaling, transport and catabolism in the presence of exogenously applied sugars (Fig. 5B,C). These expression data suggest that ATM1 is required for auxin homeostasis in seedlings, which is well-known to be required for RAM size and function. The diminished expression pattern of *DR5:GFP* in *atm1* (Fig. 6A,B) mimics that of auxin biosynthesis mutants with impaired auxin response (Brumos et al., 2018; Stepanova et al., 2008), indicating that ATM1 may be linked to auxin pathways in a sugar dependent fashion because *DR5:GFP* was increased in *atm1-1* upon sugar supplementation. While Class XI myosins have bene implicated in auxin responses this is the first report linking a Class VIII myosin to auxin-mediated root development (Abu-Abied et al., 2018; Han et al., 2021; Holweg, 2007; Holweg et al., 2003; Ojangu et al., 2018).

Previous studies have shown that cell proliferation in the root meristem are controlled by cell cycle events governed by transcription factors (Gutierrez, 2022; Li et al., 2017; Ornelas-Ayala et al., 2020; Sozzani et al., 2010; Xiong et al., 2013). Cell cycle progression from one phase to another is a tightly-regulated process and requires mitogenic signals mainly at the G1/S and G2/M transitions (Desvoyes et al., 2020; Gutierrez, 2022). Sugars are potent mitogenic signals that mediate the progression of cell cycle from G1 to S phase (Li et al., 2017; Xiong et al., 2013). Here, using EdU staining we provide novel evidence on the role of ATM1 in the activation of S-phase of the cell cycle during meristem formation in response to sugar cues (Fig. 7A,B). Similarly, we showed that the expression of H3.1-mCherry (an early S-phase marker in the PlaCCI line previously described by (Desvoyes et al., 2020)) was altered in *atm1-1* root meristems, indicating that ATM1 is a positive regulator of cell cycle progression in the RAM (Fig. 7C,D). In addition, we observed that two B-type and one P-type *CYCLIN* (*CYC*) genes are elevated in *atm1-1* in response to sucrose treatment: *CYCB1;2, CYCB1;4, CYCB1;2*. B-type cyclins control microtubule organizations during cell division in Arabidopsis, with *CYCB1;2* having established roles in root cytoskeletal regulation (Romeiro Motta et al., 2022). Furthermore, a drop in *CYC B* expression levels is associated with promotion of cell cycle progression. Thus, *atm1-1* roots appear to have impaired sucrose-induced cell cycle progression as a result of persistent cyclin expression.

Altogether, these leads us to propose a working model (Fig. 8) whereby that ATM1 positively influences RAM cell proliferation in a sugar dependent manner. In addition, ATM1 is required for normal columella differentiation. Both of these developmental processes occur in response to two key growth cues, sugars and auxin. Loss of *ATM1* impairs root morphogenesis under low light or -sugar growth conditions, suggesting that sugar signaling and/or transport may be linked to the activity of this particular myosin. In conclusion, our findings on the role of ATM1 in modulating root meristem cell cycle state and stem cell differentiation provide a missing link on the role of plant-specific Class VIII myosins in plant growth and development.

**Fig. 8.**
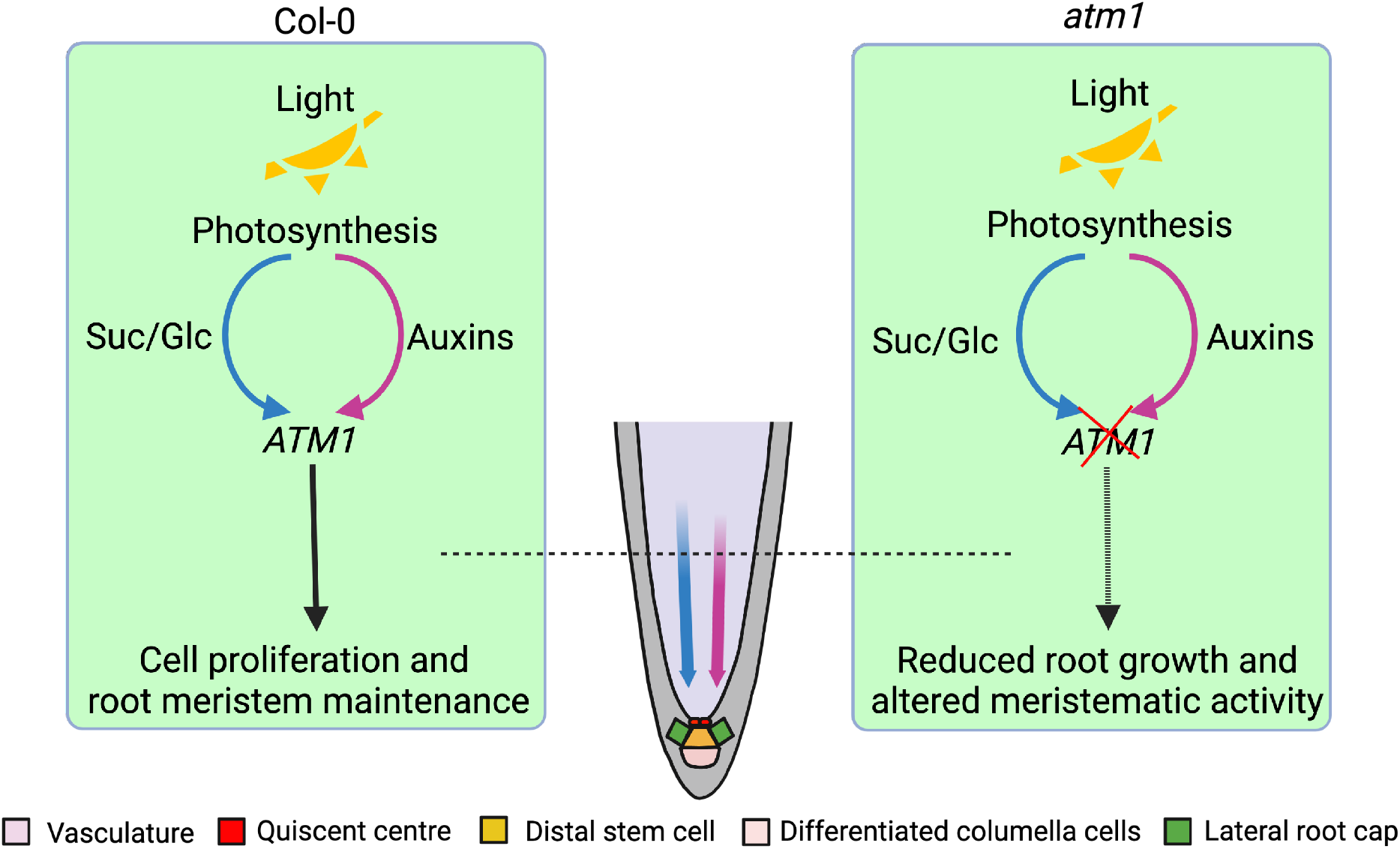
Proposed framework for ATM1 function. Wild-type root growth is stimulated by two shoot produced growth cues, sucrose and auxin. In the absence of *ATM1*, root stem cells fail to proliferate and the columella is impaired. ATM1 protein abundance is positively regulated by auxin and *atm1-1* growth defects can be partially rescued by exogenous glucose or sucrose application.

## Materials and Methods

### Plant materials and growth conditions

All *Arabidopsis thaliana* (Arabidopsis) plants used in this work were all in the Columbia (Col-0) accession background. T-DNA insertion mutant SAIL_405_B08 (*atm1-1*) was obtained from the Arabidopsis Biological Resource Center (ABRC; https://abrc.osu.edu/) and homozygous lines were identified by PCR using gene specific primers (Table S2); this null allele has been described previously (Olatunji and Kelley, 2020). The fluorescent reporter lines used in this study were crossed into *atm1-1*, carried through the F3 generation and verified by PCR based genotyping at each generation. This includes *WOX5:GFP* (Brady et al., 2007), *PET111:GFP* (Brady et al., 2007), *DR5:GFP* (Friml et al., 2003) and the Plant Cell Cycle Indicator (PlaCCI) line (Desvoyes et al., 2020).

Prior to planting, seeds were surfaced sterilized using 50% bleach and 0.01% Triton X-100 for 10 min and then washed five times with sterile water. Seeds were then imbibed in sterile water for 2 days at 4°C. The growth conditions used for experiments were 12-h white light/12-h dark, at 22°C. The light intensity was 45 μmol m-1 s-1 except otherwise stated.

For sugar-induced reactivation of root growth assays, sterilized seeds were incubated in sugar-free liquid medium half-strength (0.5X) Murashige and Skoog medium (MS) without vitamins (MSP01 Caisson), pH 5.7, for 2 days at 4°C, and then germinated in low light (15 μmol m-1 s-1, 12-h light/12-h dark, 22°C) for 3 days to enter a mitotically quiescent state as described previously (Li et al., 2019). The quiescent seedlings were then transferred to 0.5X MS medium supplemented with 15mM glucose or 15mM sucrose and incubated for 3 additional days in weak light (15 μmol m-1 s-1, 12-h light/12-h dark, 22°C) prior to root meristem and root growth analysis.

All primers used for cloning are listed in Table S2. To generate *ATM1pro::GFP-ATM1* and *ATM1pro::GUS-ATM1* constructs, a 4.5 kb genomic region upstream of the first annotated ATG in the *ATM1* gene was amplified using gene specific primers and ligated into pENTR5’ (Plasmid 27320; Addgene) to create plasmid pDO#22 (pENTR5’-ATM1pro (4.5kb)). The Green Fluorescent Protein (GFP) sequence without the stop codon was amplified from pEGAD / CD3-389 (Cutler et al., 2000) and cloned into the pDONR221 entry vector to create pDO#06 (L1-EGFP-L2). The β-glucuronidase (GUS) coding sequence was amplified from pMDC163 and cloned into entry clones pDONOR 221 and pDONOR P2R-P3 to create pDO#48 (L1-GUS-L2) and pDO#49 (R2-GUS-L3) respectively. To create the *ATM1pro::NLS-GFP-GUS* (pDO#86) construct, pEN-L1-NF-L2 (entry vector containing NLS-GFP sequences was obtained from VIB Ghent University) and pDO#49 (R2-GUS-L3) were recombined into pK7m34GW via LR Clonase reaction. A full-length *ATM1* genomic fragment (6950bp) was amplified from the first start codon to the stop codon using gene specific primers and inserted into pDONOR P2R-P3 entry vector via a BP Clonase reaction to create pDO#25 (R2-ATM1 FL-L3). The final constructs were created by cloning the entry clones into pK7m34GW destination vector using Multisite Gateway system to generate the expression clones: pDO#28 (*ATM1pro:GFP-ATM1*) and pDO#44 (*ATM1pro:GUS-ATM1*) which were individually transformed into chemically competent *Agrobacterium tumefaciens* strain GV3101 and used for the transformation of Arabidopsis plants by the floral dip method as previously described (Clough and Bent, 1998).

### Lugol Staining

Five-day old seedlings germinated on 0.5X MS plates containing 15 mM glucose or 15 mM sucrose were incubated in a root cap fixative solution (5% [w/v] formaldehyde, 5% [v/v] acetic acid, and 25% [v/v] ethanol) for 24 h and then briefly stained with Lugol solution for 30 sec. Fixed and stained roots were mounted onto microscopic slides and cleared with chloral hydrate:glycerol:water (8:3:1 ratio) before imaging with the Thunder Imaging Systems Microscope (Leica).

### 5-Ethynyl-2′-deoxyuridine (EdU) staining

EdU staining was carried out as described (Xiong et al., 2013). Briefly, surface sterilized seeds were imbibed in sugar-free liquid medium (0.5X MS without vitamins pH 5.7, for 2 days at 4°C and subsequently grown under weak light intensity (15 μmol m-1 s-1, 12-h light/12-h dark, 22°C) for 3 days in a Percival growth chamber to enter mitotically quiescent state. The quiescent seedlings were transferred into 0.5X MS medium supplemented with 15 mM glucose or 15mM sucrose for 1 day prior to EdU staining. Seedlings were stained with 1 μM EdU for 30 min and then fixed in for 30 min in 3.7% formaldehyde solution in PBS solution with 0.1% Triton X-100. After the fixative removal, the seedlings were washed three times for 10 minutes with PBS solution. The washed seedlings were then incubated in EdU detection cocktail (Invitrogen) for 30 min at room temperature in the dark, followed by PBS solution washing three times (10 min each). The root meristems of fixed seedlings were observed with a Zeiss LSM 700 confocal microscope.

### Transcriptomic and GO enrichment analyses

Total RNA was extracted from 5-day old Col-0 and *atm1-1* seedlings grown on 0.5X MS media supplemented with 15 mM glucose, 15 mM sucrose, or no sugar (“mock”); three biological replicates per genotype and treatment were collected. Total RNA was extracted using Trizol reagent followed by column clean up with a Zymo Direct-zol RNA purification kit. RNA quality was measured using a Bioanalyzer. Total RNA concentrations were determined using a NanoDrop and Qubit. QuantSeq 3′ mRNA libraries were prepared using the Lexogen 3′ mRNA-seq FWD kit and sequenced on an Illumina HiSeq 3000 as 50 bp reads. QuantSeq reads were mapped to the TAIR10 genome and differential gene expression analysis was performed using PoissonSeq implemented in R (Li et al., 2019). Raw QuantSeq data are deposited at GEO with accession GSE200917 and reviewer token qnwpuaearhsdpgt. Transcripts with a *q*-value ≤ 0.1 were assigned as differentially expressed. Gene ontology (GO) enrichment analysis was performed in PANTHER using the Arabidopsis thaliana reference genome with a Fisher’s Exact test type and a false discovery rate correction and GO terms with corrected p-value less than 0.05 were considered significantly enriched. GO terms enrichment for biological process was conducted using REVIGO (Supek et al., 2011) with a stringent dispensability cut-off (*p*-value < 0.05) and were plotted with an R script (Bonnot et al., 2019).

### Reverse Transcriptase PCR (RT-PCR)

All primers used for PCR assays are listed in Supplemental Table S2. Total RNA extracted from 5-day old whole seedlings was purified as described above. 1 μg of total RNA was used for cDNA synthesis using the LunaScript RT SuperMix Kit (New England Biolabs) according to the manufacturer’s instruction. For RT-PCR, gene-specific primers were used to amplify the target sequences for 35 cycles. PCR products were separated on 2% agarose gel.

### Confocal microscopy

Zeiss Laser Scanning Microscopy (LSM) 700 was used for confocal imaging. For confocal imaging of the root apical meristem, roots were stained with either propidium iodide (PI) or EdU. Fluorescent signals were excited with the following laser lines: GFP (488nm), and PI (555nm). The signals were then collected with the following emissions: GFP (555nm) and PI (640nm). Zeiss LSM 780 was used for imaging of *HTR3pro::HTR3-mCherry* expression from the Plant Cell Cycle Indicator (PlaCCI) transgene. The mCherry fluorescence excitation and emission used in this study was 610nm and 580-610 respectively.

### Statistical analysis

GraphPad Prism Software, version 9.3.1 (GraphPad, San Diego, CA, USA) was used for statistical analysis. The experiments were performed at least in duplicates, but only the data from one representative experiment is shown

## Data Availability

The data that support the findings of this study are openly available in NCBI GEO with accession GSE200917. All non-digital data (germplasm, plasmids and primers) are available from the corresponding author, DRK, upon reasonable request.

## Acknowledgments

This work was supported by start-up funds from Iowa State University (ISU) and an American Association of University Women Research Publication Grant to DRK. We wish to thank Cristanto Gutierrez (Centro de Biologia Molecular Severo Ochoa, CSIC-UAM, Cantoblanco, Spain) for providing the Plant Cell Cycle Indicator (PlaCCI) seeds and Bastiaan Bargmann (Virginia Tech University) for providing us with *DR5:GFP, WOX5:GFP, PET111:GFP* and *WER:GFP* seed stocks. Many thanks to Margaret Carter at the ISU Roy J. Carver High Resolution Microscopy Facility for confocal imaging assistance during the COVID19 pandemic.

## Author Contributions

DO and DRK designed the research; DO performed the research; DO and NMC analyzed data; DO and DRK wrote and edited the manuscript with input from NMC.

## References

Abu-Abied, M., Belausov, E., Hagay, S., Peremyslov, V., Dolja, V. and Sadot, E. (2018). Myosin XI-K is involved in root organogenesis, polar auxin transport, and cell division. J. Exp. Bot. 69, 2869–2881.

Bonnot, T., Gillard, M. B. and Nagel, D. H. (2019). A simple protocol for informative visualization of enriched gene ontology terms. Bio-protocol e3429–e3429.

Brady, S. M., Orlando, D. A., Lee, J.-Y., Wang, J. Y., Koch, J., Dinneny, J. R., Mace, D., Ohler, U. and Benfey, P. N. (2007). A high-resolution root spatiotemporal map reveals dominant expression patterns. Science (80-.). 318, 801–806.

Brumos, J., Robles, L. M., Yun, J., Vu, T. C., Jackson, S., Alonso, J. M. and Stepanova, A. N. (2018). Local Auxin Biosynthesis Is a Key Regulator of Plant Development. Dev. Cell 47, 306–318.e5.

Burian, A., De Reuille, P. B. and Kuhlemeier, C. (2016). Patterns of stem cell divisions contribute to plant longevity. Curr. Biol. 26, 1385–1394.

Casimiro, I., Marchant, A., Bhalerao, R. P., Beeckman, T., Dhooge, S., Swarup, R., Graham, N., Inzé, D., Sandberg, G. and Casero, P. J. (2001). Auxin transport promotes Arabidopsis lateral root initiation. Plant Cell 13, 843–852.

Casimiro, I., Beeckman, T., Graham, N., Bhalerao, R., Zhang, H., Casero, P., Sandberg, G. and Bennett, M. J. (2003). Dissecting Arabidopsis lateral root development. Trends Plant Sci. 8, 165–171.

Celenza, jr J. L., Grisafi, P. L. and Fink, G. R. (1995). A pathway for lateral root formation in Arabidopsis thaliana. Genes Dev. 9, 2131–2142.

Clark, N. M., Buckner, E., Fisher, A. P., Nelson, E. C., Nguyen, T. T., Simmons, A. R., de Luis Balaguer, M. A., Butler-Smith, T., Sheldon, P. J., Bergmann, D. C., et al. (2019). Stem-cell-ubiquitous genes spatiotemporally coordinate division through regulation of stem-cell-specific gene networks. Nat. Commun. 10, 5574.

Clough, S. J. and Bent, A. F. (1998). Floral dip: a simplified method for Agrobacterium-mediated transformation of Arabidopsis thaliana. plant J. 16, 735–743.

Cnodder, T. De, Verbelen, J.-P. and Vissenberg, K. (2006). The control of cell size and rate of elongation in the Arabidopsis root. In The expanding cell, pp. 249–269. Springer.

Cortes, S., Gromova, M., Evrard, A., Roby, C., Heyraud, A., Rolin, D. B., Raymond, P. and Brouquisse, R. M. (2003). In plants, 3-O-methylglucose is phosphorylated by hexokinase but not perceived as a sugar. Plant Physiol. 131, 824–837.

Cutler, S. R., Ehrhardt, D. W., Griffitts, J. S. and Somerville, C. R. (2000). Random GFP:: cDNA fusions enable visualization of subcellular structures in cells of Arabidopsis at a high frequency. Proc. Natl. Acad. Sci. 97, 3718–3723.

De Smet, I. and Beeckman, T. (2011). Asymmetric cell division in land plants and algae: the driving force for differentiation. Nat. Rev. Mol. Cell Biol. 12, 177–188.

Desvoyes, B., Arana-Echarri, A., Barea, M. D. and Gutierrez, C. (2020). A comprehensive fluorescent sensor for spatiotemporal cell cycle analysis in Arabidopsis. Nat. Plants 6, 1330–1334.

Ding, Z. and Friml, J. (2010). Auxin regulates distal stem cell differentiation in Arabidopsis roots. Proc. Natl. Acad. Sci. 107, 12046–12051.

Dong, P., Xiong, F., Que, Y., Wang, K., Yu, L., Li, Z. and Maozhi, R. (2015). Expression profiling and functional analysis reveals that TOR is a key player in regulating photosynthesis and phytohormone signaling pathways in Arabidopsis. Front. Plant Sci. 6, 677.

Echevarría, C., Gutierrez, C. and Desvoyes, B. (2021). Tools for Assessing Cell-Cycle Progression in Plants. Plant Cell Physiol. 62, 1231–1238.

Fernie, A. R., Roessner, U. and Geigenberger, P. (2001). The sucrose analog palatinose leads to a stimulation of sucrose degradation and starch synthesis when supplied to discs of growing potato tubers. Plant Physiol. 125, 1967–1977.

Friml, J., Vieten, A., Sauer, M., Weijers, D., Schwarz, H., Hamann, T., Offringa, R. and Jürgens, G. (2003). Efflux-dependent auxin gradients establish the apical–basal axis of Arabidopsis. Nature 426, 147–153.

Golomb, L., Abu-Abied, M., Belausov, E. and Sadot, E. (2008). Different subcellular localizations and functions of Arabidopsis myosin VIII. BMC Plant Biol. 8, 1–13.

Gonzali, S., Novi, G., Loreti, E., Paolicchi, F., Poggi, A., Alpi, A. and Perata, P. (2005). A turanose-insensitive mutant suggests a role for WOX5 in auxin homeostasis in Arabidopsis thaliana. Plant J. 44, 633–645.

Gutierrez, C. (2022). A Journey to the Core of the Plant Cell Cycle. Int. J. Mol. Sci. 23, 8154.

Han, H., Verstraeten, I., Roosjen, M., Mazur, E., Rýdza, N., Hajný, J., Ötvös, K., Weijers, D. and Friml, J. (2021). Rapid auxin-mediated phosphorylation of Myosin regulates trafficking and polarity in Arabidopsis. bioRxiv.

Haraguchi, T., Tominaga, M., Matsumoto, R., Sato, K., Nakano, A., Yamamoto, K. and Ito, K. (2014). Molecular characterization and subcellular localization of Arabidopsis class VIII myosin, ATM1. J. Biol. Chem. 289, 12343–12355.

Haraguchi, T., Duan, Z., Tamanaha, M., Ito, K. and Tominaga, M. (2019). Diversity of Plant Actin-Myosin Systems BT - The Cytoskeleton: Diverse Roles in a Plant’s Life. In (ed. Sahi, V. P.) and Baluška, F.), pp. 49–61. Cham: Springer International Publishing.

Harashima, H. and Schnittger, A. (2010). The integration of cell division, growth and differentiation. Curr. Opin. Plant Biol. 13, 66–74.

Holweg, C. L. (2007). Living markers for actin block myosin-dependent motility of plant organelles and auxin. Cell Motil. 64, 69–81.

Holweg, C., Honsel, A. and Nick, P. (2003). A myosin inhibitor impairs auxin-induced cell division. Protoplasma 222, 193–204.

Hong, J. H., Chu, H., Zhang, C., Ghosh, D., Gong, X. and Xu, J. (2015). A quantitative analysis of stem cell homeostasis in the Arabidopsis columella root cap. Front. Plant Sci. 6, 206.

Hulskamp, M., Schnittger, A. and Folkers, U. (1998). Pattern formation and cell differentiation: trichomes in Arabidopsis as a genetic model system. Int. Rev. Cytol. 186, 147–178.

Kaur, H., Manna, M., Thakur, T., Gautam, V. and Salvi, P. (2021). Imperative role of sugar signaling and transport during drought stress responses in plants. Physiol. Plant. 171, 833–848.

Kelley, D. R., Shen, Z., Walley, J. W., Chapman, E. J., Briggs, S. P. and Estelle, M. (2017). Quantitative proteomic analysis of auxin signaling during seedling development. bioRxiv 211532.

Knight, A. E. and Kendrick-Jones, J. (1993). A myosin-like protein from a higher plant. J. Mol. Biol. 231, 148–154.

Kurth, E. G., Peremyslov, V. V, Turner, H. L., Makarova, K. S., Iranzo, J., Mekhedov, S. L., Koonin, E. V and Dolja, V. V (2017). Myosin-driven transport network in plants. Proc. Natl. Acad. Sci. 114, E1385–E1394.

Lavenus, J., Goh, T., Roberts, I., Guyomarc’h, S., Lucas, M., De Smet, I., Fukaki, H., Beeckman, T., Bennett, M. and Laplaze, L. (2013). Lateral root development in Arabidopsis: fifty shades of auxin. Trends Plant Sci. 18, 450–458.

Li, L. and Sheen, J. (2016). Dynamic and diverse sugar signaling. Curr. Opin. Plant Biol. 33, 116–125.

Li, X., Cai, W., Liu, Y., Li, H., Fu, L., Liu, Z., Xu, L., Liu, H., Xu, T. and Xiong, Y. (2017). Differential TOR activation and cell proliferation in Arabidopsis root and shoot apexes. Proc. Natl. Acad. Sci. 114, 2765–2770.

Li, B., Wang, Y., Zhang, Y., Tian, W., Chong, K., Jang, J.-C. and Wang, L. (2019). PRR5, 7 and 9 positively modulate TOR signaling-mediated root cell proliferation by repressing TANDEM ZINC FINGER 1 in Arabidopsis. Nucleic Acids Res. 47, 5001–5015.

Lv, B., Yan, Z., Tian, H., Zhang, X. and Ding, Z. (2019). Local auxin biosynthesis mediates plant growth and development. Trends Plant Sci. 24, 6–9.

Mähönen, A. P., Tusscher, K. ten, Siligato, R., Smetana, O., Díaz-Triviño, S., Salojärvi, J., Wachsman, G., Prasad, K., Heidstra, R. and Scheres, B. (2014). PLETHORA gradient formation mechanism separates auxin responses. Nature 515, 125–129.

Marshall, W. F., Young, K. D., Swaffer, M., Wood, E., Nurse, P., Kimura, A., Frankel, J., Wallingford, J., Walbot, V. and Qu, X. (2012). What determines cell size? BMC Biol. 10, 1–22.

Mishra, B. S., Singh, M., Aggrawal, P. and Laxmi, A. (2009). Glucose and auxin signaling interaction in controlling Arabidopsis thaliana seedlings root growth and development. PLoS One 4, e4502.

Mishra, B. S., Sharma, M. and Laxmi, A. (2022). Role of sugar and auxin crosstalk in plant growth and development. Physiol. Plant. 174, e13546.

Nawy, T., Lee, J.-Y., Colinas, J., Wang, J. Y., Thongrod, S. C., Malamy, J. E., Birnbaum, K. and Benfey, P. N. (2005). Transcriptional Profile of the Arabidopsis Root Quiescent Center. Plant Cell 17, 1908–1925.

Nebenführ, A. and Dixit, R. (2018). Kinesins and myosins: molecular motors that coordinate cellular functions in plants. Annu. Rev. Plant Biol. 69, 329.

Ojangu, E.-L., Ilau, B., Tanner, K., Talts, K., Ihoma, E., Dolja, V. V, Paves, H. and Truve, E. (2018). Class XI Myosins Contribute to Auxin Response and Senescence-Induced Cell Death in Arabidopsis. Front. Plant Sci. 9,.

Olatunji, D. and Kelley, D. R. (2020). A role for Arabidopsis myosins in sugar-induced hypocotyl elongation. microPublication Biol. 2020, 10.17912/micropub.biology.000276.

Olatunji, D., Geelen, D. and Verstraeten, I. (2017). Control of Endogenous Auxin Levels in Plant Root Development. Int. J. Mol. Sci. 18,.

Ornelas-Ayala, D., Vega-León, R., Petrone-Mendoza, E., Garay-Arroyo, A., García-Ponce, B., Álvarez-Buylla, E. R. and Sanchez, M. de la P. (2020). ULTRAPETALA1 maintains Arabidopsis root stem cell niche independently of ARABIDOPSIS TRITHORAX1. New Phytol. 225, 1261–1272.

Park, E. and Nebenführ, A. (2013). Myosin XIK of Arabidopsis thaliana Accumulates at the Root Hair Tip and Is Required for Fast Root Hair Growth. PLoS One 8, e76745.

Peremyslov, V. V, Prokhnevsky, A. I., Avisar, D. and Dolja, V. V (2008). Two class XI myosins function in organelle trafficking and root hair development in Arabidopsis. Plant Physiol. 146, 1109–1116.

Peremyslov, V. V, Prokhnevsky, A. I. and Dolja, V. V (2010). Class XI Myosins Are Required for Development, Cell Expansion, and F-Actin Organization in Arabidopsis. Plant Cell 22, 1883 LP–1897.

Péret, B., De Rybel, B., Casimiro, I., Benková, E., Swarup, R., Laplaze, L., Beeckman, T. and Bennett, M. J. (2009). Arabidopsis lateral root development: an emerging story. Trends Plant Sci. 14, 399–408.

Perico, C. and Sparkes, I. (2018). Plant organelle dynamics: cytoskeletal control and membrane contact sites. New Phytol. 220, 381–394.

Pignocchi, C., Ivakov, A., Feil, R., Trick, M., Pike, M., Wang, T. L., Lunn, J. E. and Smith, A. M. (2021). Restriction of cytosolic sucrose hydrolysis profoundly alters development, metabolism, and gene expression in Arabidopsis roots. J. Exp. Bot. 72, 1850–1863.

Polymenis, M. and Aramayo, R. (2015). Translate to divide: control of the cell cycle by protein synthesis. Microb. cell (Graz, Austria) 2, 94–104.

Racolta, A., Bryan, A. C. and Tax, F. E. (2014). The receptor-like kinases GSO1 and GSO2 together regulate root growth in Arabidopsis through control of cell division and cell fate specification. Dev. Dyn. 243, 257–278.

Ramon, M., Rolland, F. and Sheen, J. (2008). Sugar sensing and signaling. Arab. B. 6, e0117–e0117.

Reddy, A. S. and Day, I. S. (2001). Analysis of the myosins encoded in the recently completed Arabidopsis thaliana genome sequence. Genome Biol. 2,.

Reichelt, S., Knight, A. E., Hodge, T. P., Baluska, F., Samaj, J., Volkmann, D. and Kendrick-Jones, J. (1999). Characterization of the unconventional myosin VIII in plant cells and its localization at the post-cytokinetic cell wall. Plant J. 19, 555–567.

Rolland, F., Baena-Gonzalez, E. and Sheen, J. (2006). Sugar sensing and signaling in plants: conserved and novel mechanisms. Annu. Rev. Plant Biol. 57, 675–709.

Romeiro Motta, M., Zhao, X., Pastuglia, M., Belcram, K., Roodbarkelari, F., Komaki, M., Harashima, H., Komaki, S., Kumar, M. and Bulankova, P. (2022). B1-type cyclins control microtubule organization during cell division in Arabidopsis. EMBO Rep. 23, e53995.

Roychoudhry, S. and Kepinski, S. (2022). Auxin in root development. Cold Spring Harb. Perspect. Biol. 14, a039933.

Ryan, J. M. and Nebenführ, A. (2018). Update on Myosin Motors: Molecular Mechanisms and Physiological Functions. Plant Physiol. 176, 119 LP–127.

Sairanen, I., Novák, O., Pěnčík, A., Ikeda, Y., Jones, B., Sandberg, G. and Ljung, K. (2012). Soluble Carbohydrates Regulate Auxin Biosynthesis via PIF Proteins in Arabidopsis. Plant Cell 24, 4907–4916.

Santner, A., Calderon-Villalobos, L. I. A. and Estelle, M. (2009). Plant hormones are versatile chemical regulators of plant growth. Nat. Chem. Biol. 5, 301–307.

Santuari, L., Sanchez-Perez, G. F., Luijten, M., Rutjens, B., Terpstra, I., Berke, L., Gorte, M., Prasad, K., Bao, D. and Timmermans-Hereijgers, J. L. P. M. (2016). The PLETHORA gene regulatory network guides growth and cell differentiation in Arabidopsis roots. Plant Cell 28, 2937–2951.

Shi, B. and Vernoux, T. (2022). Hormonal control of cell identity and growth in the shoot apical meristem. Curr. Opin. Plant Biol. 65, 102111.

Shishkova, S., Rost, T. L. and Dubrovsky, J. G. (2008). Determinate root growth and meristem maintenance in angiosperms. Ann. Bot. 101, 319–340.

Shulse, C. N., Cole, B. J., Ciobanu, D., Lin, J., Yoshinaga, Y., Gouran, M., Turco, G. M., Zhu, Y., O’Malley, R. C. and Brady, S. M. (2019). High-throughput single-cell transcriptome profiling of plant cell types. Cell Rep. 27, 2241–2247.

Singh, S., Yadav, S., Singh, A., Mahima, M., Singh, A., Gautam, V. and Sarkar, A. K. (2020). Auxin signaling modulates LATERAL ROOT PRIMORDIUM 1 (LRP 1) expression during lateral root development in Arabidopsis. Plant J. 101, 87–100.

Sozzani, R., Cui, H., Moreno-Risueno, M. A., Busch, W., Van Norman, J. M., Vernoux, T., Brady, S. M., Dewitte, W., Murray, J. A. H. and Benfey, P. N. (2010). Spatiotemporal regulation of cell-cycle genes by SHORTROOT links patterning and growth. Nature 466, 128–132.

Stals, H. and Inzé, D. (2001). When plant cells decide to divide. Trends Plant Sci. 6, 359–364.

Stepanova, A. N., Robertson-Hoyt, J., Yun, J., Benavente, L. M., Xie, D.-Y., Dolezal, K., Schlereth, A., Jurgens, G. and Alonso, J. M. (2008). TAA1-mediated auxin biosynthesis is essential for hormone crosstalk and plant development. Cell 133, 177–191.

Su, S.-H., Gibbs, N. M., Jancewicz, A. L. and Masson, P. H. (2017). Molecular Mechanisms of Root Gravitropism. Curr. Biol. 27, R964–R972.

Supek, F., Bošnjak, M., Škunca, N. and Šmuc, T. (2011). REVIGO summarizes and visualizes long lists of gene ontology terms. PLoS One 6, e21800.

Szövényi, P., Waller, M. and Kirbis, A. (2019). Evolution of the plant body plan. Curr. Top. Dev. Biol. 131, 1–34.

Titus, M. A. (2018). Myosin-driven intracellular transport. Cold Spring Harb. Perspect. Biol. 10, a021972.

Torres-Martínez, H. H., Napsucialy-Mendivil, S. and Dubrovsky, J. G. (2022). Cellular and molecular bases of lateral root initiation and morphogenesis. Curr. Opin. Plant Biol. 65, 102115.

Wang, J., Song, J., Clark, G. and Roux, S. J. (2018). ANN1 and ANN2 Function in Post-Phloem Sugar Transport in Root Tips to Affect Primary Root Growth. Plant Physiol. 178, 390–401.

Xiong, Y., McCormack, M., Li, L., Hall, Q., Xiang, C. and Sheen, J. (2013). Glucose–TOR signalling reprograms the transcriptome and activates meristems. Nature 496, 181–186.

Zluhan-Martínez, E., López-Ruíz, B. A., García-Gómez, M. L., García-Ponce, B., de la Paz Sánchez, M., Álvarez-Buylla, E. R. and Garay-Arroyo, A. (2021). Integrative Roles of Phytohormones on Cell Proliferation, Elongation and Differentiation in the Arabidopsis thaliana Primary Root. Front. Plant Sci. 12,.

