## Supplementary material for "The Class VIII myosin ATM1 is required for root apical meristem function": Figure S1

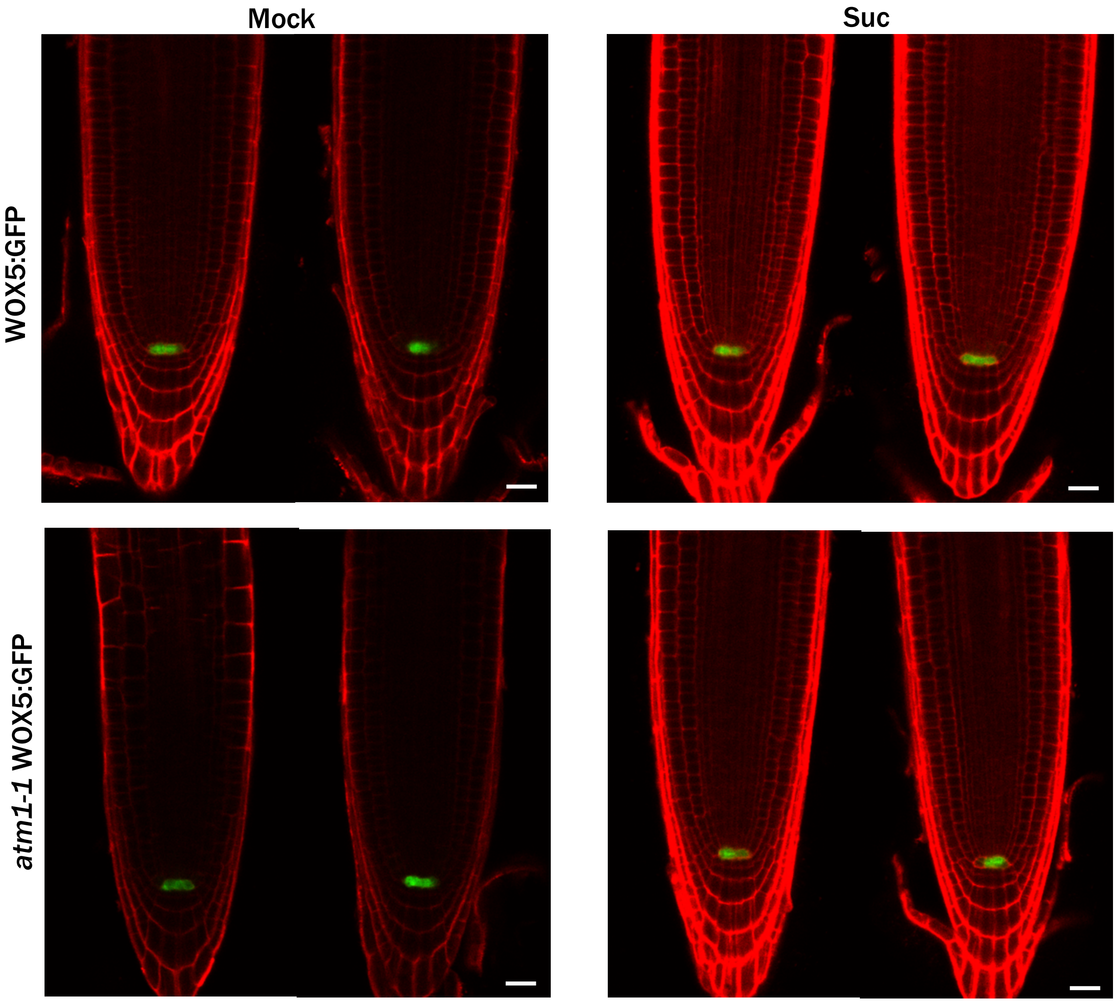


**Fig. S1. The expression of the quiescent marker WOX: GFP is normal in *atm1-1*.** The roots of 5-day-old seedlings grown on 0.5X MS medium (mock) or supplemented with 15 mM sucrose were counterstained with propidium iodide. Bars = 20 μm.
